# Safety margin evaluations for transtympanic laser stimulation of the cochlea from the outer ear in Mongolian gerbils (*Meriones unguiculatus*)

**DOI:** 10.1101/2022.10.31.514500

**Authors:** Aya Okamoto, Miku Uenaka, Yuki Ito, Tomohiro Miyasaka, Koji Toda, Shizuko Hiryu, Kohta I. Kobayasi, Yuta Tamai

**Affiliations:** Neuroethology and Bioengineering, Graduate School of Life and Medical Sciences, Doshisha University, Kyotanabe, Kyoto, Japan; Neuropathology, Graduate School of Life and Medical Sciences, Doshisha University, Kyotanabe, Kyoto, Japan; Department of Psychology, Keio University, Tokyo, Japan

**Keywords:** Auditory prosthesis, Infrared laser stimulation, Auditory perception, Head-fixed classical conditioning

## Abstract

Infrared laser cochlear stimulation has been suggested as an alternative to traditional auditory prostheses. Previous research has mainly examined the short-term effects of laser stimulation, but to effectively use this technique, it is necessary to study whether prolonged laser exposure can safely and sustainably induce cochlear responses. This study assessed the effect of laser-induced cochlear damage using classically-conditioned Mongolian gerbils. The white noise was presented as a conditioned stimulus, and reward licking was recorded as a conditioned response. After training, the subject’s cochlea was exposed to a continuous pulsed laser for 15 h. The licking rate drastically decreased after the exposure of 26.4 W/cm^2^, while it did not change after 6.6 W/cm^2^ or weaker. These findings showed, for the first time, that the safety margin of long-term, at least several hours, cochlear laser stimulation exists and will contribute to the safe and effective delimitation of laser parameters in future research.

## Introduction

Hearing impairment is a prevalent sensory disorder worldwide. Approximately 466 million people, 5% of the world’s population, suffer from hearing deficits, resulting in severe communication difficulties and social isolation ^1–3^. Hearing aids are among the most widely used therapeutic options to compensate for these problems. These devices improve speech comprehension and social interaction in hard-of-hearing individuals ^4–6^. However, these patients require an intact middle ear to ensure sufficient and proper hearing. Hard-of-hearing individuals with middle ear dysfunction must use an implantable auditory prosthesis (e.g., bone-anchored hearing devices, middle ear implants, or cochlear implants) that bypasses the function of the middle ear to stimulate the peripheral auditory system. One of the most significant drawbacks preventing the widespread use of these devices is surgical intervention, which increases the risk of complications post-surgery ^7–9^.

Infrared laser stimulation has been discussed as a possible alternative to conventional auditory devices ^10^. The advantage of laser stimulation is that it can activate spatially focused cell populations in a non-contact manner without the need to deliver any exogenous agent into tissues, such as a viral vector ^11^. Early research by Izzo *et al.* ^12^ suggested that pulsed laser stimulation of the auditory nerves induced a cochlear response. Several later studies that focused on the detailed relationship between the irradiation site of the cochlea and laser-evoked neural response indicated that laser stimulation could elicit spatially specific neural activities ^13–16^. That is, precisely manipulating the laser stimulation system can improve the spectral resolution of cochlear implants. Our previous studies leveraged the benefit of the contactless feature of laser stimulation to reduce the invasiveness of surgical implantation of the auditory prosthesis. They revealed that transtympanic laser stimulation from the outer ear could evoke a cochlear response, bypassing the middle ear function ^17, 18^. The auditory cortical activity elicited by transtympanic laser stimulation depends on stimulus intensity ^19^. These results imply that a hearing prosthesis with infrared laser stimulation could be a noninvasive alternative to the conventional implantable auditory prosthesis.

For the clinical application of transtympanic laser stimulation to auditory prostheses, the safety margin of laser irradiation should be assessed. Because the laser-evoked neural response is probably related to the thermal elevation of the tissue ^20–22^, thermal build-up might be the main reason for laser-induced damage ^23–25^. Therefore, thermal damage to the peripheral auditory system can occur when a laser with high radiant energy is continuously presented. Some studies have investigated laser irradiation trauma in the cochlea ^12, 17, 26, 27^. Goyal *et al.* ^28^ compared the relationship between radiant energy in continuous laser exposure and the reduction of the laser-evoked electrophysiological response in the cochlea, reporting that the injury threshold for laser stimulation at 250 Hz was between 25 and 30 µJ/pulse. However, no study has described the influence of laser-induced thermal elevation on cochlear tissues or the effect of thermal physiological damage on auditory perception.

A comprehensive evaluation of laser irradiation trauma elucidates the effects of laser-induced damage on auditory perception. It provides a precise picture of the effect of laser stimulation on the auditory peripheral system. It assessed laser-induced damage characteristics of the peripheral auditory system by comparing electrophysiological, histological, and behavioral alterations following continuous laser exposure. Here, we performed (1) histological examination of spiral ganglions with hematoxylin and eosin (HE) staining, (2) electrophysiological recording of the laser-evoked cochlear response during laser irradiation, and (3) head-fixed classical conditioning with auditory stimulus (see Fig. 1 for details).

**Fig. 1.**
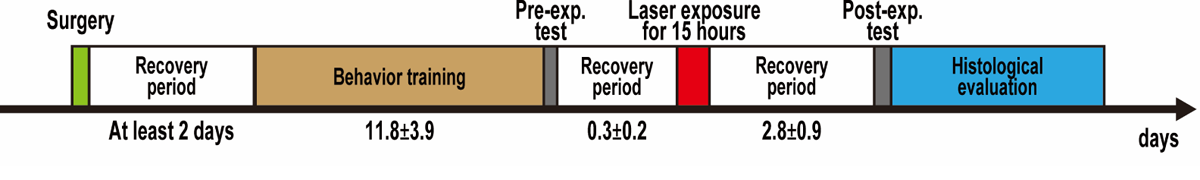
Time course of the experiment. Before head-fixed classical conditioning, surgery was performed to set the head plate and recording electrode on the subjects and to impair hearing sensitivity in the right ear. At least two days after the surgery, the training session was conducted to associate the relationship between the auditory cue and the liquid reward for 11.8 ± 3.9 days (Mean ± SEM, n=18). The pre-exposure test was conducted after the training was completed. Subsequently, continuous laser exposure was presented for 15 h after recovery (0.3 ± 0.2 days). The cochlear response was recorded during the laser exposure. The post-exposure test was carried out 2.8±0.9 (Mean±SEM) days after the laser exposure. Finally, the subjects were euthanized on the same day as the post-exposure test, and histological inspection was performed.

This study performed a comprehensive safety evaluation of laser irradiation trauma in Mongolian gerbils (*Meriones unguiculatus*). The Mongolian gerbil is a standard animal model for investigating auditory physiology ^e.g., 29–31^. Several studies have used gerbils to investigate the mechanism of human speech perception because gerbils have a higher sensitivity to the frequency range of human speech, including harmonics (250–3000 Hz), than other rodents ^32–35^. These studies suggest that the Mongolian gerbils is an excellent animal model for studying the mechanism of human hearing. Furthermore, the anatomical structure of the cochlea in gerbils is relatively large and shares several similarities with humans compared to other rodents ^36, 37^. This anatomical similarity allows us to evaluate the properties of laser-induced thermal distribution in the cochlea, which is essential data for the clinical application of laser stimulation in hearing prostheses. Therefore, the safety evaluation of transtympanic laser stimulation in Mongolian gerbils will help to establish safe and effective laser stimulation parameters.

## Results

### Continuous laser irradiation of 26.4 W/cm^2^ or more increased the cochlear surface temperature

Before the histological examination, the surface temperature of the cochlea was recorded to investigate the thermal elevation following continuous laser irradiation (Fig. 2). The surface temperatures did not increase from 24 ℃ (i.e., room temperature) using the 6.6 W/cm^2^ laser irradiation or less; the laser 26.4 W/cm^2^ or more increased the surface temperatures immediately within 1 min into laser irradiation. The temperature rise reached a plateau approximately 5 min after the laser onset (Fig. 2A). Average surface temperature at 10 min after the laser irradiation onset was described in Fig. 2B and showed that the increasing temperature induced by 26.4, 52.8, and 105.6 W/cm^2^ lasers were 31.6, 36.6, and 44.7 ℃, respectively. The ANOVA revealed that the effect of radiant energy was statistically significant (F(5,17)=21.26, P<0.001, one-way ANOVA). Fig. 2C shows the lateral and medial surface temperatures induced by laser irradiation at 105.6 W/cm^2^ on the lateral side of the cochlea. The average temperatures of the lateral and medial sides resulted in approximately 41.2 and 32.5 ℃, respectively, and the temperature difference between the left and the right sides was statistically significant (P<0.01, paired t-test). The medial side temperature was approximately in line with that of the lateral side irradiated by the laser at 26.4–52.8 W/cm^2^.

**Fig. 2.**
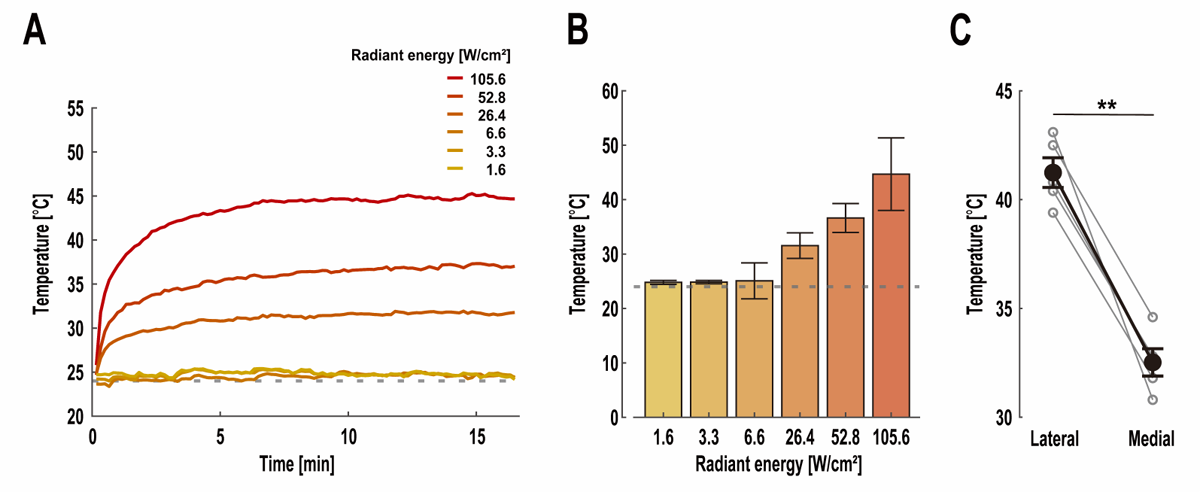
Thermal elevation of the cochlear surface caused by laser irradiation. (A) Time course of the average temperature rise on the irradiation site of the cochlear surface. (B) Surface temperature at 10 min after the laser onset in (A). Error bars indicate the standard deviation (1.6 W/cm^2^: n = 3, others: n = 4). (C) Temperature difference between the lateral and medial sides of the cochlear surface following the laser irradiation of 105.6 W/cm^2^ (n=5). Glay open circles show the temperature in each cochlea. In (A) and (B), the gray dotted line shows the room temperature (24 ℃). Error bars describe the standard error of the mean. **: *P* < 0.01.

### Continuous laser irradiation mainly damaged the spiral ganglion neuron of the apical-middle turn on the lateral side

Horizontal sections of the cochlea were subjected to HE staining to evaluate the effect of laser exposure on the spiral ganglion neuron density (Fig. 3A). Lower spiral ganglion neuron density was found in the cochlea continuously irradiated by the laser of 105.6 W/cm^2^ than in less than 6.6 W/cm^2^ (Fig. 3B). The middle turn of the cochlea irradiated by the laser at 105.6 W/cm^2^ is depicted in Fig. 3C. A comparison of the lateral and medial sides showed a remarkable decrease in the cell density on the lateral side, mainly irradiated by the transtympanic laser, compared to the medial side. On the lateral side of the cochlea, the spiral ganglion density difference between the left and right ears significantly decreased (r=-0.87, P<0.05, Pearson’s correlation analysis) as radiant energy intensified, whereas on the medial side, it did not (r=-0.45, P=0.23, Pearson’s correlation analysis) (Fig. 3D). The cell density decrement on the medial side after laser exposure of 105.6 W/cm^2^ approximately coincides with that on the lateral side irradiated by the laser with 26.4–52.8 W/cm^2^. Fig. 3E shows the spiral ganglion neuron density difference on the lateral side of the apical-middle and basal turn of the cochlea. The apical-middle turn of the cell density in the left ear was significantly lower than that in the right ear (P<0.05, one-sample t-test). In contrast, the basal turn of the cell density was less affected than the apical-middle side (P=0.19, one-sample t-test) (Fig. 3E).

**Fig. 3.**
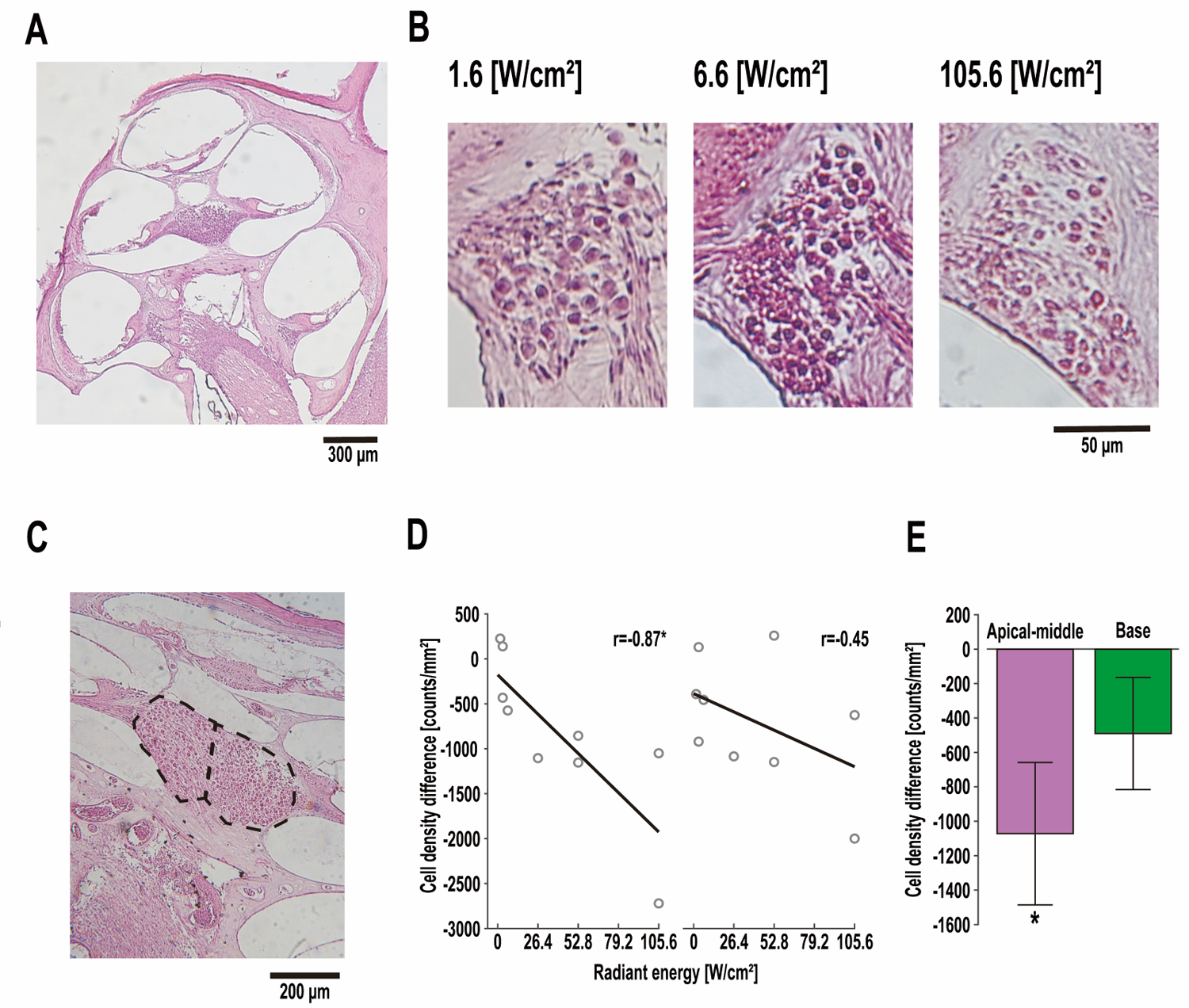
Laser-induced lesion in spiral ganglion neurons. (A) Horizontal section of laser irradiated cochlea showing all turns (radiant energy: 6.6 W/cm^2^). (B) Density of spiral ganglion neurons in the lateral side of the basal turn after laser exposure of 1.6 W/cm^2^ (left), 6.6 W/cm^2^ (middle), and 105.6 W/cm^2^ (right). (C) Lateral (left) and medial (right) sides of the spiral ganglion neuron density in middle turn. The dotted line separates the lateral and medial sides of the spiral ganglion area in the middle turn. (D) Change in the cell density after laser exposure on lateral (left) and medial (right) sides of the cochlea. The cell density difference was defined as the mean spiral ganglion density difference between the laser irradiated (left ear) and the opposite (right ear) cochlea. Gray open circles indicate individual data (n=9). Correlation coefficients were calculated using Pearson’s correlation analysis. (E) Mean cell density difference between the laser irradiated and opposite cochlea in apical-middle (left) and basal (right) turn on the lateral side. Data from all radiant energy conditions were pooled in the analysis. *: *P* < 0.05.

### The cochlear response with the laser of 26.4 W/cm^2^ or more was reduced during the continuous laser exposure

Fig. 4 shows the cochlear response during continuous laser exposure for 15 h. The manifest damage in cochlear response was not observed by laser exposure up to 6.6 W/cm^2^, while the cochlear response elicited by laser exposure of 105.6 W/cm^2^ decreased (Fig. 4A). The time course of the relative response amplitude showed that continuous laser exposure at 105.6 W/cm^2^ caused a drastic decrease in the laser-evoked response amplitude within ca. 65 min (Fig. 4B). In comparison, laser irradiation of 6.6 W/cm^2^ or less did not. Fig. 4C shows the relationship between the laser irradiation time and percentage of response amplitude. Laser irradiation over 26.4 W/cm^2^ drastically increased the average amplitude decrement, while laser stimulation with up to 26.4 W/cm^2^ did not (Fig. 4D).

**Fig. 4.**
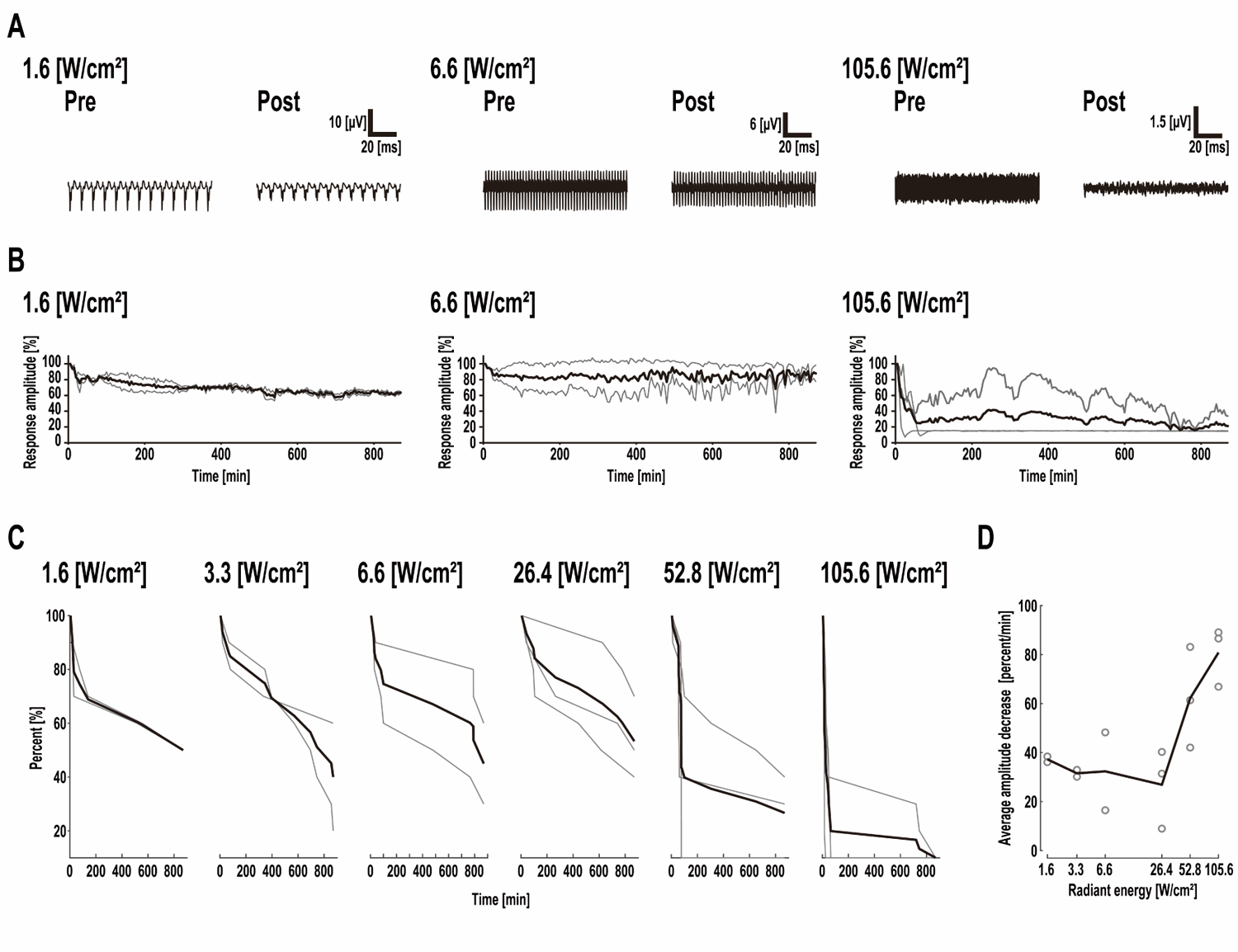
Cochlear function assessment during continuous laser exposure. (A) Cochlear response during continuous pulsed laser irradiation of 1.6 (left), 6.6 (middle), and 105.6 (right) W/cm^2^ (i.e., 125, 500, and 8000 pulses/s). (B) Time course of the cochlear response amplitude elicited by the 1.6 (left), 6.6 (middle), and 105.6 (right) W/cm^2^ laser stimulation. (C) Relationship between laser irradiation time and percent of the response amplitude. (D) Average amplitude decrease caused by continuous laser irradiation depending on the radiant energy. Gray open circles indicate individual data (n=15). Gray lines in (B) and (C) show individual data.

### The continuous laser exposure within 6.6 W/cm^2^ did not change the behavioral level, while the 26.4 W/cm2 or more laser did

Fig. 5A shows the licking behavior evoked by white noise (80 dB SPL) before and after 1.6, 6.6, and 105.6 W/cm^2^ laser exposure. The laser exposure at 1.6 and 6.6 W/cm^2^ did not cause an observable behavioral response change, while an apparent decrement of the licking rate was observed after 105.6 W/cm^2^ laser exposure. The behavioral changes after laser exposure are depicted in Fig. 5B. Laser irradiation of the cochlea over 6.6 W/cm^2^ drastically changed the behavioral level (Fig. 5B). It caused the delta behavioral level (DBL) to reach out of the safety margin, which was calculated using three standard deviations of DBL in the control condition (Fig. 5C).

**Fig. 5.**
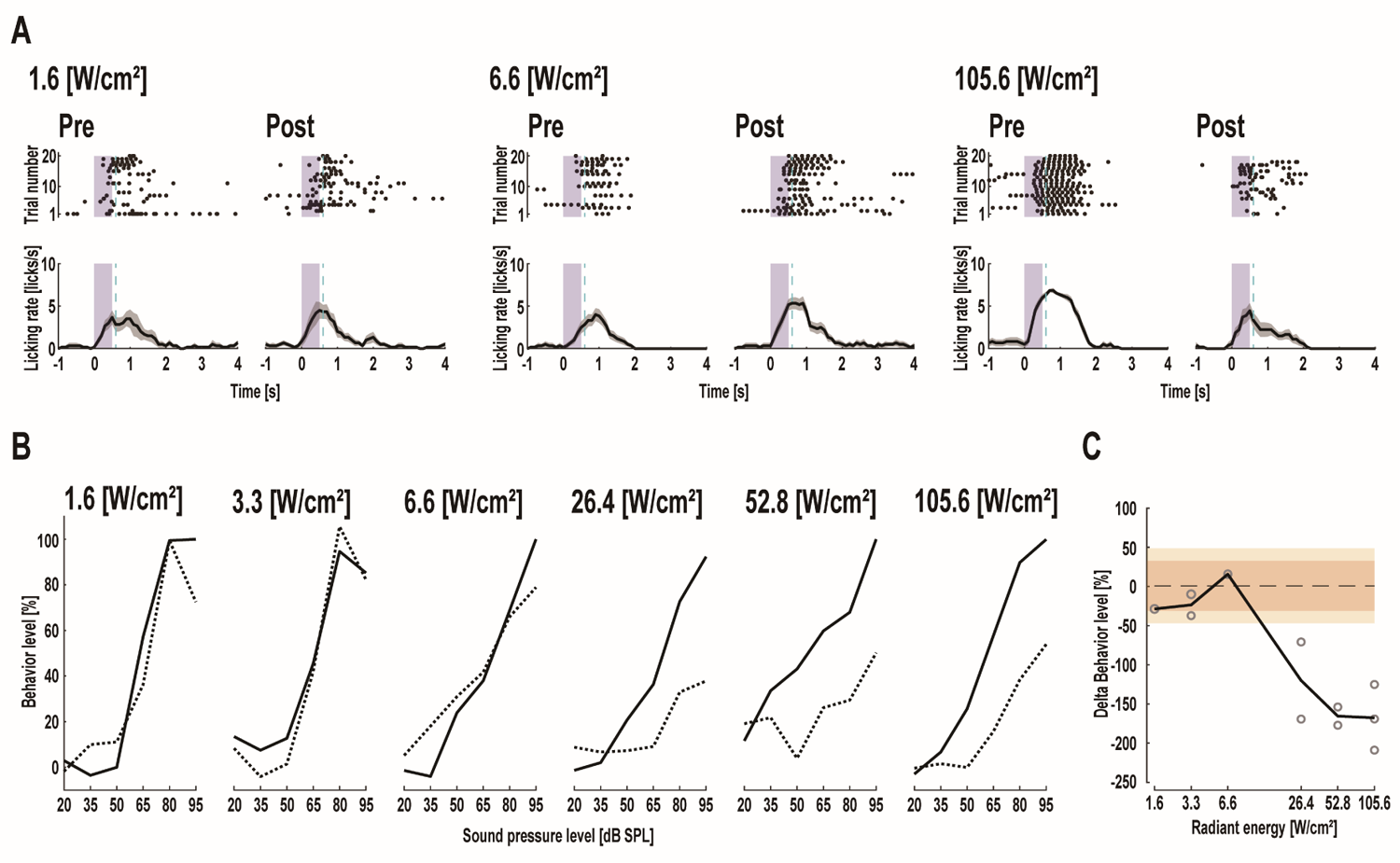
Influence of continuous laser irradiation on the auditory-evoked behavioral response. (A) Auditory-evoked licking behavior pre- and post-laser exposure of 1.6 (left), 6.6 (middle), and 105.6 (right) W/cm^2^. Individual trials and mean peri-stimulus time histograms (PSTH) of the licking behavior elicited by auditory stimuli (80 dB SPL) are shown in top and bottom figures, respectively. The blue zones indicate auditory stimulation periods. The light blue dotted lines show the reward timing in the training trials. Black plots in the top figure show licking timing. (B) Behavioral level alternation after continuous laser irradiation. Solid and dotted lines show pre- and post-behavioral levels, respectively. The behavioral level was determined as the ratio of delta licking rate over the maximum delta licking rate of the pre-laser exposure (1.6 W/cm^2^: n=1, 3.3 W/cm^2^: n=2, 6.6 W/cm^2^: n=2, 26.4 W/cm^2^: n=2, 52.8 W/cm^2^: n=2, 105.6 W/cm^2^: n=3). (C) Change in the delta behavioral level between pre- and post-laser exposure. In the control condition, delta behavioral level was measured by comparing the pre- and post-placing the subjects on the experimental setup for 15 h (n=9). Gray dotted line shows the mean of the delta behavioral level in the control condition, and colored areas indicate two (red) and three (orange) standard deviations. The safety margin was determined as the mean ± three standard deviations (i.e., orange area). Gray open cycles represent delta behavioral response in each subject (n = 18).

## Discussion

This study demonstrated that the laser stimulation with a higher average power density triggered the thermal elevation of the cochlear surface (Fig. 2). Because the conventional mechanism of the laser-evoked cochlear response is the optothermal effect, which causes an excessive temperature increase in the cells ^20–22^, several studies indicated that laser-induced thermal elevation and possible thermal-damage on the cochlea might constitute a significant limitation in applying the laser to the auditory prosthesis ^24, 26–28, 38^. Our study demonstrated that cochlear surface temperature significantly increased as radiant energy increased (Fig. 2AB). The surface temperature continued to rise for approximately 5 min after laser onset, and then the elevated temperature reached a plateau (Fig. 2A). These results suggest that continuous laser irradiation (minute order) elevates cochlear temperature compared with short-term irradiation. The continuous laser irradiation at 105.6 W/cm^2^ caused an average surface temperature up to 44.7 °C, potentially causing thermal tissue damage ^39^. These results demonstrate that continuous laser stimulation bears the risk of thermal damage to the cochlear tissues when a laser with high radiant energy is used.

A comparison of the thermal elevation of the cochlear surface with histological damage of spiral ganglion neurons revealed that the thermal distribution of the cochlear surface affected the laser irradiation trauma of the spiral ganglion neurons. Our previous study showed that transtympanic lasers mainly irradiated the middle turn of the lateral side of the cochlea ^18^. This study implies that the highest peak of the thermal distribution of the cochlear surface might be observed in the primary irradiated area, and the temperature could decrease from the lateral to the medial side. It also showed that the thermal elevation of the lateral side caused by laser irradiation of 105.6 W/cm^2^ was significantly higher than that on the medial side. The 105.6 W/cm^2^ laser-induced temperature rise of the medial side reached 32.5 ℃, which agreed with the lateral side temperature caused by the laser at 26.4–52.8 W/cm^2^ (Fig. 2C). These results suggest that approximately half of the radiant energy of laser irradiation reached the medial side of the cochlea and agreed with the laser irradiation site in our previous study ^18^. In addition, a more significant decrease in cell density was observed on the lateral side of the apical-middle turn than on the basal turn (Fig. 3E). These results coincide with the radiant energy distribution (i.e., thermal distribution) of the cochlear surface and suggest that the spatial injury profile of ganglion neurons depends on the thermal distribution of the cochlear surface.

The electrophysiological cochlear response was recorded to evaluate the acute damage threshold of continuous laser stimulation. This response is widely used as a sensitive probe for physiological cochlear function ^40–42^. Previous studies, including ours ^12, 17, 26^, have assessed laser-irradiation trauma of the cochlea by recording the cochlear response induced by continuous laser stimulation. These studies suggest that the laser safely evokes cochlear responses without causing severe irradiation damage. However, Goyal *et al*. ^28^ reported that continuous laser irradiation with higher radiant energy induced thermal damage to the cochlear function. This indicates that the injury threshold of laser exposure is between 25 and 30 µJ/pulse at 250 pulses/s (i.e., approximately 47.1 and 56.5 W/cm^2^). Our results showed that the cochlea kept responding to laser stimulation of 26.4 W/cm^2^ or less, while the laser at 52.8 W/cm^2^ or above remarkably decreased the cochlear response (Fig. 4D). This result suggests that the electrophysiological damage threshold of the laser stimulation is between 26.4 and 52.8 W/cm^2^, and the injury threshold is in line with the previous study by Goyal *et al*. ^28^.

Measuring the impact of thermal-induced physiological deterioration in the cochlea on auditory perception is crucial. Behavioral assessment of hearing loss generally provides compelling evidence of auditory perceptual consequences following compromised auditory processing. Therefore, evaluating auditory perception following thermal injury in the cochlea is an essential step toward the clinical application of laser stimulation to auditory prostheses. This study showed that the auditory-evoked licking behavior did not change after laser exposure of 6.6 W/cm^2^ or weaker, but drastically decreased after 26.4 W/cm^2^ or above. These results suggest that the thermal injury threshold for transtympanic laser stimulation is between 6.6 and 26.4 W/cm^2^. Therefore, transtympanic laser stimulation over 6.6 W/cm^2^ could damage the subject’s peripheral auditory system and be out of the safety margin.

A comparison of the behavioral and electrophysiological results revealed that the thermal damage threshold for the auditory-evoked behavior alteration was slightly lower than that for the laser-evoked cochlear response. Here, the laser irradiates the tympanic membrane, auditory ossicles, and cochlear tissues from the outer ear to induce the cochlear response ^18^; laser stimulation bears the risk of lesioning more peripheral auditory function from the origin of the laser-evoked response. Our previous study revealed that laser irradiation (λ=1871 nm) from the outer ear bypasses the tympanic membrane and auditory ossicles because these tissues do not have higher absorbance to the infrared laser^18^. However, some studies investigated the effect of the laser irradiation of the tympanic membrane and auditory ossicles in *vivo* ^43^ and in *vitro* ^44^. This indicates that the laser irradiation of 27.9 W/cm^2^ or above at 532 nm could cause thermal damage. Although these results are not directly comparable to our data because of the difference in wavelength-dependent absorbance in the tympanic membrane and auditory ossicles, as some studies described ^45–47^, thermal damage to the tympanic membrane and auditory ossicles might be one reason why continuous laser exposure affects the auditory-evoked response more than the laser-evoked response.

In addition to the tympanic membrane and auditory ossicles, hair cells in the cochlea can be damaged by laser irradiation. The physiological mechanism of the laser-evoked cochlear response is still debated whether the response is hair-cell-mediated ^11^. Hence, whether the laser-evoked response here characterized the state of cochlear function is unclear, including that of hair cells. To reveal the physiological mechanism of laser stimulation in the cochlea, some studies ^48–50^ compared laser-evoked auditory responses in normal hearing and chemically compromised cochlea and reported that laser-evoked auditory responses were only observed in the normal hearing cochlea. These studies suggest that the cochlear response elicited by laser stimulation is mediated by hair cells.

In contrast, Richter *et al*. ^51^ revealed that 30–40 dB chemical deafening of the auditory threshold did not affect the threshold of the laser-evoked cochlear response. Tan *et al*. ^52^ demonstrated that laser stimulation could elicit auditory neural activity in congenitally deaf mice in which the cochlea could not transmit synapses between the inner hair cells and spiral ganglion neurons.

In addition, when animals have an impaired hearing threshold, auditory masking does not affect laser-induced responses ^53^. These results suggest that laser irradiation of the cochlea directly activates the auditory nerves, bypassing hair cells. If the laser activates the cochlear nerves without needing hair cell depolarization, the laser-evoked response does not characterize the state of the hair cells. Therefore, our results show that the auditory-evoked response might be more vulnerable to thermal injury than the laser-evoked response (Fig. 4 and 5). The effect of thermal damage to hair cells on laser-evoked cochlear responses must be considered in the future.

In summary, we assessed the laser-induced damage characteristics of the peripheral auditory system through histological, electrophysiological, and behavioral tests. Histological evaluation of the cochlea after continuous laser exposure revealed that the laser-induced thermal distribution of the cochlear surface coincided with the spatial injury profile of spiral ganglion neurons. The measurement of auditory-evoked behavioral and laser-evoked electrophysiological responses indicate that cochlear response during the laser exposure of 26.4 W/cm^2^ or weaker did not decrease. In contrast, auditory-induced behavioral response drastically decreased after 26.4 W/cm^2^ or higher. This discrepancy in the injury threshold suggests that laser stimulation bears the risk of lesioning more peripheral auditory functions from the origin of the laser-evoked response. Evaluating perceptual consequences indicated that the thermal injury threshold for transtympanic laser stimulation in Mongolian gerbils is between 6.6 and 26.4 W/cm^2^. Therefore, transtympanic laser under 6.6 W/cm^2^ could be available without concern for acute injury. A comprehensive safety margin assessment is an essential step toward the practical application of transtympanic laser stimulation in auditory prostheses.

### Limitations of the Study

One of the limitations of this research is that the safety of chronic transtympanic laser irradiation of the cochlea was not investigated. The present study showed that the transtympanic laser at a power of 6.6 W/cm^2^ or weaker did not cause acute damage. However, hearing prostheses can be expected to be used over a long period of time, albeit intermittently rather than continuously. In order for transtympanic laser stimulation to be used in clinical applications, further studies on investigating the long-term safety of auditory perception needs to be conducted.

Another limitation is that safety on the human peripheral auditory organs has not been evaluated. Thermal damage from laser irradiation depends on the laser absorbance and tissue heat dissipation, which is related to the size and composition of the peripheral auditory system. Our previous study revealed that thicker tympanic membranes have higher absorption; human tympanic membranes thicker those of gerbils might be at greater risk for thermal damage ^18^. As for the cochlea, the size of the cochlea is more massive in humans than in gerbils ^37^. Goyal et al. ^28^ discussed that the larger cochlea has a higher rate at which heat is distributed and dissipated, suggesting that the energy threshold for damage may be higher in humans than in small animals. Therefore, the safety margin of the peripheral auditory system in humans might be different from our current results. For practical application of transtympanic laser stimulation, a more detailed evaluation of laser absorption and heat dissipation in the human peripheral auditory system needs to be performed in the future.

## Materials and methods

### Animals

All experimental procedures were performed according to the guidelines established by the Ethics Review Committee of Doshisha University. The experimental protocols were approved by the Animal Experimental Committee of Doshisha University. A total of 33 experimentally naive Mongolian gerbils (fourteen females, nineteen males; *Meriones unguiculatus*) aged 3–18 months were used here. All gerbils were bred and reared in a laboratory. Each animal was housed with 2–5 gerbils in a 20 (W) × 40 (L) × 17 cm (H) cage, with free access to food and water. The animal room had a 12-h light-dark schedule, a temperature of 22–23 °C, and relative humidity of approximately 50%. All subjects were randomly assigned to each experimental group.

### Animal surgery

Mongolian gerbils were deeply anesthetized with ketamine (47 mg/kg i.m.) and xylazine (9.3 mg/kg i.m.). Maintenance doses of ketamine (17.5 mg/kg) and xylazine (7.0 mg/kg) were injected every 30–50 min or if the animals showed signs of increasing arousal (e.g., voluntary whisker movement). The body temperature of the gerbils was maintained using heating pads placed beneath the animals. The left side of the bulla was exposed by incision of the muscle and skin from the shoulder to the jaw, and a hole (diameter: 1 mm) was made on the bulla. A silver electrode (Nilaco, Tokyo, Japan; diameter: 0.13 mm; impedance < 20 kΩ) was inserted into the hole and hooked onto the bony rim of the round window to record cochlear responses. The electrode was stabilized with a bulla using acrylic glue and dental cement (Provinice; Shofu, Kyoto, Japan). The left pinna was removed to provide a clear view of the tympanic membrane. A liquid of acrylic glue and dental cement (Provinice; Shofu, Kyoto, Japan) was mounted into the right ear for approximately 50 dB SPL attenuation of the sound pressure reaching the right ear.

The parietal and temporal sides of the fur and the skin of the subject’s head were shaved. The temporal muscle was removed after applying a local anesthetic (xylocaine gel; Aspen Japan, Tokyo, Japan). The skull was carefully cleaned with 0.3% sodium hypochlorite solution and saline. A small hole was made in the dorsal skull 2 mm anterior to the bregma and 1 mm lateral to the midline, and a reference electrode was inserted into the hole. A V-shaped metal plate (F-911; Hilogik, Osaka, Japan) was attached to the skull using acrylic dental resin (Super bond C & B; Sun Medical, Shiga, Japan) and dental cement (Provinice; Shofu, Kyoto, Japan), allowing the subjects to be head-fixed during the experiment. All subjects were singly housed during recovery for at least two days before training began.

### Auditory stimuli

White noise (700–44,800 Hz) was used as an auditory stimulus, with a duration of 500 ms. The auditory stimulus was presented via a doom tweeter (FT28D; Fostex, Tokyo, Japan) from 15 cm above the subject’s head. The stimulus was generated using a digital-to-analog converter (Octa-Capture; Roland, Shizuoka, Japan) with a sampling rate of 192 kHz. The signal was intensified using an amplifier (A-10; Pioneer, Tokyo, Japan). The sound pressure level was calibrated to 20, 35, 50, 65, 80, and 95 dB SPL using a microphone (Type 7016; Aco, Tokyo, Japan).

### Apparatus and procedure in the behavioral experiment

Behavioral experiments were conducted in a soundproof box 60 (W) × 60 (L) × 60 cm (H). Licking recordings in head-fixed animals were performed as previously described^54–57^. Each subject was positioned on a covered elevated platform (custom-designed and 3D printed). A drinking spout was placed 1 mm in front of the mouth of the animal, and the clamp and drinking spout positions were adjusted to detect licking behavior easily. A touch sensor (DCTS-10; Sankei Kizai, Tokyo, Japan), connected between the drinking spout and the metal sheet of the stage, recorded the timing of licks at 2000 samples/s. The recorded licking timing was stored on a computer via Arduino Uno (Arduino, Ivrea, Italy) controlled by Python 3.7 on Spyder 501. All behaviors were recorded using a video camera (HD-5000; Microsoft, Washington, USA).

After at least 2 days of recovery from surgery, the gerbils were water-restricted in their home cage. The training session typically comprised 400 trials daily. Head-fixed participants were allowed to voluntarily lick the spout at all times. A 3.2 µl drop of water (unconditioned stimulus: US) was delivered through the tube every 7–13 s after presenting an 85 dB SPL of white noise (conditioned stimulus: CS) controlled by MATLAB (MathWorks, Massachusetts, USA) and a microcontroller (Arduino Uno; Arduino, Ivrea, Italy) with a custom-made relay circuit.

After animals showed high licking rates following the auditory cue, catch trials were conducted to evaluate the subject’s perception. The test session comprised rewarded and non-rewarded trials (i.e., catch trials). Catch trials were included to assess the subjects’ perceptions without reward feedback. White noise of 20, 35, 50, 65, 80, or 95 dB SPL was presented in the catch trials. During a test session, 280 rewarded trials and 120 catch trials were conducted in a pseudo-random order. In the catch trials, each sound intensity was tested 20 times. The test session was performed pre- and post-laser exposure.

### Recording and analysis of the behavioral data

An averaged histogram of licking behavior was used to evaluate the time course of behavioral responses. The histograms were calculated in a 100 ms bin size and smoothened using a 300-ms moving average window. Peak licking rates, durations, peak latencies, and response latencies were extracted from the time course of licking behavior to assess the behavioral response. Peak licking rates and peak latencies were defined as the licking rate and time when the licking rate reached the maximum value within 4 s after stimulus onset, respectively. The baseline licking rate was defined as the pre-stimulus licking rate obtained for 1 s, and duration was quantified as the temporal length of licking behavior >50% of the licking rate from the mean baseline licking rate to the peak licking rate. Response latency was determined by measuring the time between stimulus onset and the behavioral response reaching the mean plus three standard deviations of the baseline licking rate. The delta licking rate was defined as the difference between the mean baseline licking rate and the post-stimulus licking rate to evaluate the response amplitude. The analysis range of post-stimulus licking behavior was 0.2–1.2 s because the behavioral response was mostly observed 0.2 s after stimulus onset (see Supplemental Fig. 1 for details). The behavioral level was calculated by standardizing the highest delta licking behavior in the pre-exposure test. The delta behavioral level (DBL) between the pre- and post-exposure test was defined as the summation of the difference in the behavioral level for all sound pressure levels to evaluate the effect of continuous laser exposure on auditory perception.

### Laser exposure

A repetitive pulsed infrared laser was used for laser exposure. The individual pulse of the laser had a 100 µs rectangular shape in intensity. It was generated using a diode laser stimulation system (BWF-OEM-1850; B&W TEK, Delaware, USA) with a wavelength (λ=1871) comparable to those previously reported ^12, 38, 50^. The voltage commands for the laser stimuli were generated using a function generator (WF1974; NF, Kanagawa, Japan). The repetition frequency of the pulsed laser was changed to 125, 250, 500, 2000, 4000, and 8000 Hz to adjust the amount of radiant energy reaching the cochlea to 1.6, 3.3, 6,6, 26.4, 52.8, and 105.6 W/cm^2^, respectively. The radiant energy and beam width at the cochlea (i.e., 2.2 mm from the fiber tip) were measured using a digital power meter (PM100D; Thorlabs, Tokyo, Japan) with a thermal power sensor (S302C; Thorlabs, Tokyo, Japan) and an InGaAs biased detector (DET10D/M; Thorlabs, Tokyo, Japan), respectively. Our previous study measured the laser-beam profile ^18^.

An optic fiber (diameter: 100 μm; NA, 0.22) was inserted mediolaterally into the left outer ear canal with the tip angled rostrally by 10° and dorsally by 5° using a micromanipulator (MM-3; Narishige, Tokyo, Japan) in awake head-fixated subjects. The optic fiber tip was set at c.a. 0.7 mm in front of the tympanic membrane to ensure that the cochlea was irradiated transtympanically from the outer ear as described in our previous reports ^17–19^.

Each subject’s cochlea was continuously stimulated by laser irradiation from the outer ear for 15 h. Continuous laser exposure was performed on the same covered elevated platform in the soundproof box as in the behavioral experiment. The temperature in the soundproof box was adjusted to approximately 26 ℃, and a small pee pad was placed beneath the subject to prevent the subject’s temperature from decreasing. After laser exposure and behavioral experiments, the subjects were overdosed with an intraperitoneal injection of pentobarbital (200 mg/kg). Laser-induced cochlear response was continuously recorded throughout the study period. The cochleae were histologically investigated to clarify the thermal damage caused by laser irradiation.

### Recording and analysis of the cochlear response during laser exposure

After the pre-exposure test, cochlear response was recorded in awake subjects during continuous laser exposure. The cochlear response was amplified 1000 times using a bio-amplifier (MEG-1200; Nihon Kohden, Tokyo, Japan) and stored on a computer via an analog-to-digital converter (Octa-capture; Roland, Shizuoka, Japan) at a sampling rate of 96,000 Hz.

The recorded signal was processed using a band-pass filter (100–10,000 Hz) and separated at 10 ms intervals. The cochlear signal was averaged 1000 times every 10 s for 15 h, and the response amplitude of the averaged cochlear signal was evaluated using the amplitude spectrum distribution. To obtain high-frequency resolution, 32,768 points of zero paddings were performed on the head and end of the averaged signal. The amplitude spectrum was extracted using fast Fourier transforms (FFT, 16,384 points with a Hanning window); therefore, the frequency resolution was 5.9 Hz. The peak of the amplitude spectrum at the stimulus repetition frequency was extracted as the response amplitude. The mean response amplitude was calculated at 5 min intervals standardized by the first 5 min mean amplitude.

### Preparation of histological evaluation

Following the post-exposure test session, the laser-irradiated subjects were euthanized by an overdose of pentobarbital (200 mg/kg i.p.). After postmortem perfusion with phosphate buffered saline (PBS) and 4% paraformaldehyde in PBS, the animals were decapitated and fixed in the same fixative for two days in a refrigerator at 4 ℃. Subsequently, the tissues were stored in cooled PBS containing 0.02% sodium azide.

The paraformaldehyde-fixed tissues were decalcified by immersing in a mixture solution of formic acid (0.28 g/ml), citric acid (0.1 g/ml), and sodium hydroxide (0.02 g/ml) for 7–10 days. The decalcifying solution was exchanged every 2–3 days. After the decalcification, the samples were preserved in PBS with 0.02% sodium azide.

Decalcified head tissues were dehydrated using a gradually ascending concentration of ethanol (70–100%) for 2 days followed by immersion in 100% xylene and embedding into a paraffin block. Horizontally, paraffin-embedded sections with the thickness of 6 µm was processed using a microtome (HM 430; Thermo, Tokyo, Japan).

Cochlear sections between 7 and 8 mm ventral to the sagittal midline from each subject were examined. Sections were deparaffinized by immersion in 100% xylene. After the deparaffinization, sections were hydrated with 100% ethanol and distilled water. Sections were stained with Mayer’s hematoxylin solution and rinsed with distilled water. After differentiation with 70% ethanol containing 0.1% HCL, the sections were stained with Putt’s eosin solution. HE-stained sections were re-immersed in 100% ethanol and cleared in xylene. All stained sections were mounted in mounting medium (HSR solution; Sysmex, Hyogo, Japan). Specimens were photographed using a digital camera (BX53; Olympus, Tokyo, Japan) under a light microscope (EX-F1; Casio, Tokyo, Japan). Histological analysis was performed using ImageJ software ^58, 59^. The main part of the analyzed spiral ganglions might be afferent Type-I neurons because they are 95 % of the total population of the spiral ganglion.

### Behavioral data analysis for a safety margin evaluation

Each subject was placed on a covered elevated platform for 15 h in a temperature-controlled soundproof box, similar to the laser exposure experiment without presenting laser irradiation. We compared the performance of the classical conditioning task before and after placement on the covered elevated platform. The mean and standard deviation of the control condition were measured to determine the safety margin of continuous laser irradiation. The safety margin was defined as the mean ± three standard deviations of the DBL in the control condition (n=9).

### Correlation between physiological damage in the cochlea and behavioral alternation

To determine whether these histological and functional deficits in the cochlea were associated with behavioral alterations, we correlated the change in spiral ganglion neuron density and amplitude of the cochlear response with the behavioral response. Further details are provided in Supplemental Fig. 2.

### Recording thermal elevation of the cochlear surface

Eleven gerbils were euthanized (pentobarbital; 200 mg/kg) and decapitated for cochlear surface thermal measurements. Both sides of the cochlea were extracted from the skulls under a stereomicroscope. The cochlea was placed in a Petri dish and stabilized with dental wax. The laser fiber tip was directed vertically to the surface of the cochlea and was set to 2.2 mm from the cochlear surface. Continuous laser irradiation with 1.6, 3.3, 6.6, 26.4, 52.8, and 105.6 W/cm^2^ was performed for 1 h, and the surface temperature elevations were measured using a thermal camera (FLIR C3-X; Teledyne FLIR LLC, Oregon, USA), which was positioned at 21 cm from the cochlear surface with vertically 45° angle.

To acquire the thermal elevation on the lateral and medial sides of the cochlear surface, a thermocouple (MSM-3905; Biomedica, Osaka, Japan) was used. The tip of the thermocouple sensor was in contact with the second turn of the cochlear surface on one side for 1 h. After recording the temperature on one side, the specimen was left for at least 15 min to cool down to room temperature (∼24 °C), and then the temperature on another side was measured.

### Flow of the behavioral, electrophysiological, and histological procedure

Surgery, behavior training, pre-exposure test, laser exposure for 15 h, post-exposure test, and histological evaluation were performed in this order. The recovery period was performed after surgery before and after laser exposure. Further details are provided in Fig. 1.

## Acknowledgement

The authors would like to thank Hiroshi Riquimaroux for his support during the early stages of this work. We also thank Kazuki Tanaka, Mizuki Katayama, and Keito Hosokawa for technical support and Editage (www.editage.com) for English language editing. This research was financially supported by the Sasakawa Scientific Research Grant No. 2022–6027 (A.O.), the Japan Society for the Promotion of Science (JSPS) KAKENHI Grants Nos. 22H02946 (T.M.), 21H03469 (K.I.K.), 21K21322 (Y.T.), the JSPS Overseas Research Fellowship (Y.T.), and Keio Academic Development Fund (K.T.).

## Author contribution

Conceptualization: A.O., KI.K., and Y.T.; Data curation: A.O. and M.U.; Formal analysis: A.O. and Y.T.; Funding acquisition: A.O., T.M., K.T., KI.K., and Y.T.; Investigation: A.O. and Y.T.; Methodology: Y.I., T.M., and K.T.; Project administration: KI.K. and Y.T.; Software: A.O. and Y.T.; Supervision: S.H.; Validation: A.O.; Visualization: A.O. and Y.T.; Writing-Original Draft: A.O. and Y.T.; Writing-Review&Editing: A.O., M.U., Y.I., T.M., K.T., S.H., KI.K., and Y.T.

## Data availability statement

All data needed to evaluate the conclusions are shown in the paper and supplemental figures. Additional data related to this paper are available from the corresponding author upon reasonable request.

## Supplemental Figures

**Supplemental Fig. 1.**
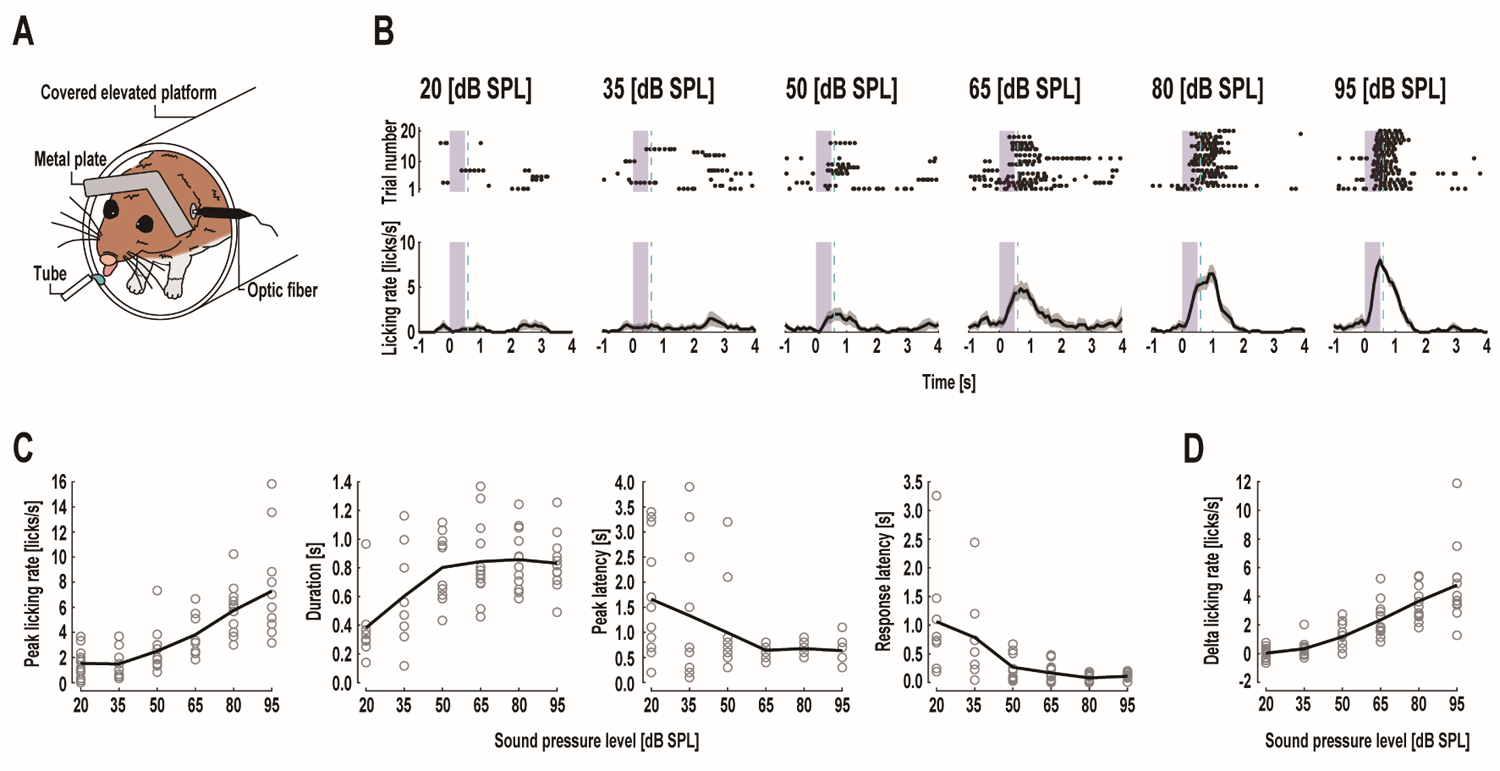
Auditory-induced licking behavior in head-fixed Mongolian gerbils and behavioral profile depending on the sound pressure level. (A) Schematic of the head-fixed setup. We investigated the perception of auditory stimuli by using head-fixed classical conditioning. A drop of water was delivered through the tube after presenting white noise, and the timing of licks was recorded using a touch sensor, as in previous studies. (B) Individual trial (top) and mean peri-stimulus time histogram (PSTH) (bottom) of the licking behavior elicited by auditory stimuli with various sound pressure levels. The blue zones indicate auditory stimulation periods. The light blue dotted lines show the reward timing in the training trials. Black plots in the top figure show licking timing. Increasing the intensity of white noise elicited a systematic change in behavioral responses. (C) Intensity dependence of the peak licking rate, duration, peak latency, and response latency. As the sound pressure level increased from 20 to 95 dB SPL, peak licking rate was increased from 1.52 to 7.28 licks/s, duration was increased from 0.38 to 0.83 s, peak latency was decreased from 1.66 to 0.63 s, response latency was decreased from 1.06 to 0.11 s. (D) Mean auditory-evoked delta licking rate. The delta licking rate was defined as the response amplitude to quantify the response amplitude of licking behavior. The delta licking rate significantly increased from 0.04 to 4.78 licks/s depending on the intensity of the white noise. In (C) and (D), the open grey cycles show the individual data (n=12).

**Supplemental Fig. 2.**
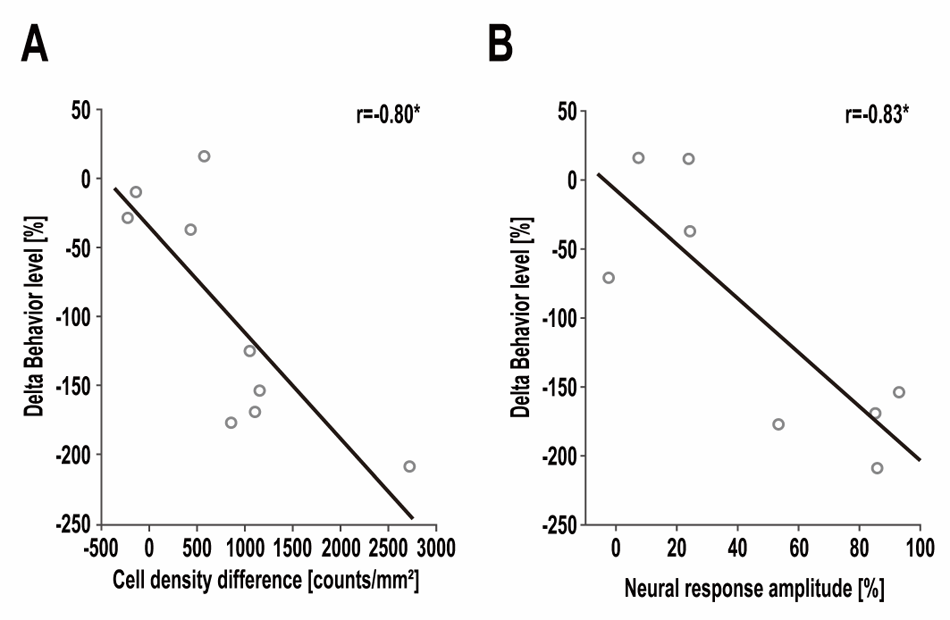
Change in the delta behavioral level depending on the decrease in the cell density difference. (A) and neural response amplitude (B). Histological and behavioral data (n=9) or electrophysiological and behavioral data (n=8) were obtained from the subjects. The correlation coefficients were calculated using Pearson’s correlation analysis. As the spiral ganglion neuron density decreased, the delta behavioral level significantly decreased (r=-0.80, P<0.05, Pearson’s correlation analysis). A similar tendency was observed in the relationship between the decrement of the cochlear response and behavioral response; the decreasing amplitude of the cochlear response significantly reduced the behavioral response (r=-0.83, P<0.05, Pearson’s correlation analysis). These results suggest that auditory perception is affected by the physiological deterioration of the cochlea. The open circles indicate individual data. *: *P* < 0.05.

